# Polymodal Sensory Thalamic Inputs to the Rat Lateral Amygdala are Facilitated by Chronic Ethanol Exposure and Regulate Withdrawal-associated Anxiety

**DOI:** 10.1101/557256

**Authors:** Melissa Morales, Molly M. McGinnis, Ann M. Chappell, Brian C. Parrish, Brian A. McCool

## Abstract

Thalamic projections to the lateral amygdala regulate the acquisition of conditioned aversive and reward-related behaviors. Recent work suggests that exposure to chronic ethanol up-regulates presynaptic function of lateral amygdala *stria terminalis* inputs which contain projections from somatosensory thalamic nuclei. To understand potential contributions by thalamic inputs and their role in the expression of withdrawal-associated aversive behaviors, we integrated optogenetic and chemogenetic approaches with *in vitro* measures of synaptic function and anxiety-like behavior. We found that expression of Channelrhodopsin in the caudal extension of the posterior thalamic group (cPO) produced monosynaptic glutamatergic synaptic responses in lateral amygdala principal neurons that could be inhibited by co-expression of the hM4-Gi-DREADD. Chronic ethanol exposure increased glutamate release from these cPO terminals but did not impact inhibition by the DREADD agonist, CNO. Systemic injection of CNO specifically reduced withdrawal-related increases in anxiety-like behaviors in animals expressing the Gi-DREADD in cPO. And, microinjection of CNO directly into the lateral amygdala mimicked this anti-anxiety effect. These findings suggest that the cPO-LA circuit is vulnerable to chronic ethanol exposure and plays an important role in regulating anxiety-like behavior following chronic ethanol exposure.

## Introduction

Alcohol use disorder (AUD) is characterized by excessive alcohol intake and a negative emotional state when alcohol is not in use. As AUD and anxiety are strongly connected and anxiety following cessation of drinking is a strong contributor to relapse, animal models are necessary to understand not only the behavioral/functional consequences of alcohol withdrawal, but also the neurophysiological and circuitry that are involved. For instance, chronic ethanol exposure via vapor inhalation induces anxiety-like behavior and increases ethanol consumption during withdrawal—two hallmarks of ethanol dependence (reviewed in (Koob, 2003; Koob, 2013)). In addition, ethanol dependence produces neurophysiological changes in several key brain regions associated with the fear/anxiety circuit, including the lateral/basolateral amygdala (McCool, 2011; McCool et al., 2010).

The basolateral amygdala is a highly conserved structure made up predominantly of glutamatergic principal neurons and a smaller proportion of GABAergic interneurons (McDonald, 1982; McDonald, 1985; McDonald, 1992). The interactions between and activity of these neuronal populations regulate the expression of anxiety-like and fear-related behaviors (Siuda et al., 2016). The basolateral amygdala consists of the lateral amygdala (LA) and basal subdivisions. Located dorsally, the LA receives sensory information from the sensory cortex and thalamus and projects extensively to the basolateral amygdala as well as distant brain regions. These projections ultimately regulate physiological/psychological responses related to reward as well as conditioned and unconditioned aversive behaviors.

Chronic ethanol exposure produces robust effects on both anxiety-like behavior and synaptic transmission onto basolateral amygdala principal neurons (Christian et al., 2013; Christian et al., 2012; Diaz et al., 2011b; Lack et al., 2007; Morales et al., 2018; Robinson et al., 2016). Importantly, chronic intermittent ethanol (CIE) vapor exposure produces distinct, pre- and post-synaptic alterations in afferents that are both input- and exposure-duration (Morales et al., 2018). These works have shown that glutamatergic afferents arriving via the *stria terminalis* (ST; dorsal to the central amygdala) express a presynaptic form of ‘ethanol plasticity’ and develop with shorter exposures compared to the postsynaptic potentiation expressed at external capsule (EC) regardless of age or sex. This suggests that the initial presynaptic facilitation of ST inputs may be critical for the development of behavioral and synaptic withdrawal symptomology. These CIE/withdrawal data parallel fear-conditioning studies that demonstrate time-dependent activation of the ST and then the EC causes use-dependent, associative postsynaptic plasticity at the cortical/lateral inputs onto basolateral amygdala neurons (Cho et al., 2012; Humeau et al., 2003). The interaction between medial presynaptic glutamate release and lateral postsynaptic potentiation may reflect a crucial mechanism for modulating fear- and ethanol-conditioned behaviors.

Research on the circuitry controlling fear and anxiety responses has recently focused on interactions between the BLA and prefrontal cortex. However, early work in this field demonstrated a critical role for thalamic structures. For example, the LA receives sensory information from several thalamic subdivisions. Notably, the caudal extension of the posterior thalamic group (cPO) – which includes the intralaminar nucleus (PIN), the triangular cell subdivision (PoT), the medial subdivision of the medial geniculate (mMG), and the suprageniculate nucleus (Sg) – is a collection of polymodal sensory areas located just medial to the medial geniculate nucleus (MG) and lateral to the anterior pretectal (APT) area. The cPO sends dense projections to the LA via the *stria terminalis* (Doron and Ledoux, 1999; LeDoux et al., 1990; Nitecka et al., 1979); and permanent lesions to the PIN, Sg, and mMG reduce elevations in c-fos immunoreactivity in the LA following exposure to foot-shocks (Lanuza et al., 2008). These studies highlight the importance of cPO inputs to the LA for fear-related behaviors; however, their precise interactions during alcohol withdrawal are unknown. Therefore, the current study utilized chemo- and optogenetic approaches with behavioral and electrophysiological methods to determine the role of cPO inputs into the LA in mediating anxiety-like behavior following ethanol withdrawal. We show that these cPO inputs are both sensitive to chronic ethanol exposure and are important modulators of withdrawal-associated anxiety.

## Methods

### Animals

Upon arrival, four to five-week old male Sprague-Dawley rats (Envigo; Indianapolis, IN) were pair-housed and maintained on a reverse 12:12 h light dark cycle (lights on at 9PM). Rats were given unlimited access to standard rat chow and water throughout the experimental procedures. All animal care procedures were in accordance with the NIH Guide for the Care and Use of Laboratory Animals in accordance with the Society for Neuroscience Policies on the Use of Animals and Humans in Neuroscience Research and approved by the Wake Forest School of Medicine Animal Care and Use Committee.

### Surgical procedures

Rats were anesthetized with isoflurane (2.5-5% for induction and maintenance) and underwent aseptic surgery using a stereotaxic apparatus (Neurostar, Germany). Bilateral injections of AAV vectors (rAAV5/CamKII-hChR2(H134R)-eYFP-WPRE and rAAV8/hSyn-hM4D-mcherry; co-injected at 1:1 ratio; both from UNC vector core) were administered using a Harvard Apparatus (Holliston, MA) pump at a rate of 0.1 μl/min, for a total volume of 1μl per side (cPO injection coordinates: −5.76 anterior/posterior; ±3.00 medial/lateral; and, −6.20 dorsal/ventral). We removed sutures 1 week later and pair-housed the animals. For electrophysiology studies and systemic clozapine-N-oxide injections, viral expression progressed for four weeks prior to the chronic intermittent ethanol vapor inhalation (below).

For the behavioral studies using BLA microinjections, rats underwent a second surgery two weeks after microinjection of the viral constructs to implant chronic guide cannulae (Plastics One; Roanoke, VA). We placed cannulae bilaterally into the BLA (BLA coordinates: −2.80 anterior/posterior; ±5.00 medial/lateral; and, −7.58 dorsal/ventral) and affixed them to the skull with dental cement according to previous publications (Diaz et al., 2011a; McCool and Chappell, 2007; McCool et al., 2014). To retain the patency of the guide cannulae, we placed sterile obturators into each cannulae and replaced them as necessary. Rats were pair-housed 3 days following this second surgery. One week prior to the CIE exposure, we microinjected all rats with 0.5 μl/side aCSF to habituate them to the procedural manipulations.

### Chronic intermittent ethanol (CIE) vapor exposure

Chronic intermittent ethanol (CIE) vapor exposure lasted for 7 days using standard procedures (Christian et al., 2012; Lack et al., 2007; Morales et al., 2018; Robinson and McCool, 2015). Briefly, we placed pair-housed rats in their home cages into a larger, custom-built Plexiglas chamber (Triad Plastics, Winston-Salem, NC). At the onset of the light cycle (9pm EST), we applied ethanol vapor (15-20 mg/L) for 12h each day into the exposure chambers. We housed ethanol-naïve control animals under identical conditions except that they received room-air only while in the chambers. This 7 day exposure is sufficient to increase glutamate release from stria terminalis inputs to the BLA using electrical stimulation (Christian et al., 2013; Morales et al., 2018). Body weights were collected daily; and, tail blood samples were collected during the CIE exposure to monitor blood ethanol concentrations and adjust ethanol vapor levels as necessary. Blood samples were collected from any individual animal only once during the exposure; and blood ethanol concentrations were determined from plasma using a commercially available alcohol dehydrogenase/NADH enzymatic assay (Diagnostic Chemicals Limited, Oxford CT). The average BEC during the CIE vapor exposure was 219.15±12.65mg/dl.

### Whole-cell Patch Clamp Electrophysiology

#### Slice preparation

Rats were anesthetized with isoflurane and decapitated. Brains were immediately removed and incubated for 5 minutes in an ice-cold sucrose-modified artificial cerebral spinal fluid (aCSF) solution containing (in mM): 180 Sucrose, 30 NaCl, 4.5 KCl, 1 MgCl_2_·6H_2_O, 26 NaHCO_3_, 1.2 NaH_2_PO_4_, 10 glucose, 0.10 ketamine. Coronal amygdala slices (400μM) were prepared using a VT1200/S vibrating blade microtome (Leica, Buffalo Grove, IL and incubated for ≥1 h in a room temperature, oxygenated, standard aCSF solution containing (in mM): 126 NaCl, 3 KCl, 1.25 NaH_2_PO_4_, 2 MgSO_4_, 26 NaHCO_3_, 10 glucose, and 2 CaCl_2_ prior to recordings. All chemicals were from Sigma-Aldrich (St. Louis, MO) or Tocris (Ellisville, Missouri).

#### Whole-cellpatch-clamp recording

We performed whole-cell recordings from lateral amygdala neurons within coronal brain slices according to established procedures (Morales et al., 2018). In a submersion-type recording chamber, room temperature aCSF continuously perfused brain slices at a rate of 2ml/min. We pharmacologically isolated glutamatergic responses with the GABA_A_ antagonist, picrotoxin (100μM in the bath aCSF). The intracellular recording solution for synaptic responses contained (in mM): 145 CsOH, 10 EGTA, 5 NaCl, 1 MgCl, 10 HEPES, 0.4 QX314, 1 CaCl_2_, 4 Mg-ATP, 0.4 Na-GTP (pH ~7.25, modified with gluconic acid, osmolarity between 285-295 mmol/kg, modified with sucrose). For current-clamp recordings, K-gluconate replaced Cs-gluconate in this solution. We collected cellular/synaptic responses using an Axopatch 700B amplifier (Molecular Devices, Foster City, CA) and later analyzed the responses off-line using pClamp software (Molecular Devices, Foster City, CA). The electrophysiological characteristics of lateral amygdala principal neurons - high membrane capacitance (>100 pF) and low access resistance (≤25 MΩ) (Rainnie et al., 1993; Washburn and Moises, 1992) – were used as inclusion criteria for analysis. Changes of >20% for whole-call capacitance or access resistance at any point during the duration of the recording excluded cells from analysis.

#### Optogenetics & Chemogenetics

To measure light-evoked action potentials or glutamate release (ChR2-mediated) from cPo terminals in the lateral amygdala, we placed a naked fiber optic cable (400μm diameter) immediately above the tissue (directly over the injection site or stria terminalis) and delivered brief light stimulations from a 473nm laser triggered by 5V TTL stimulations controlled from the acquisition software. The 473nm light durations varied from 2msec to 45msec in the cPO recordings; and, we used a 5msec stimulus delivered at 0.5Hz for the synaptic responses in the lateral amygdala. We assessed glutamate release probability with two light-evoked responses separated by a 50msec inter-stimulus interval and derived a paired-pulse ratio (PPR) from light-evoked EPSC amplitudes using a conservative estimate ([Peak 2 amplitude - Peak 1 amplitude] / Peak 1 amplitude; (Christian et al., 2013; Diaz et al., 2011b; Lack et al., 2007; Morales et al., 2018)). Gi DREADD modulation of synaptic responses was measured by acute application of the DREADD agonist, cloxapine-N-oxide (CNO, 10μM). Following a 5min baseline period, we perfused CNO onto slices for 10 min, followed by a 10 min washout period. We applied CNO to brain slices only once; CNO-inhibition only partially recovered after extensive washing.

### Elevated Plus Maze

We used previously standardized procedures to assess anxiety-like behavior during withdrawal using the Elevated Plus Maze (EPM, 5min test; (Diaz et al., 2011a; Morales et al., 2015; Morales et al., 2018)). For the systemic injection studies, we injected rats with CNO (5 mg/kg i.p.) or saline (1 ml/kg i.p) 30 min prior to placement onto the central junction of the EPM facing an open arm. For the intracerebral studies, we microinjected rats, via an injection needle which extended 1.0mm past the guide cannula, with CNO (300μM) or aCSF (both at 0.5μl/side over 1min) 5min prior to placement onto the EPM. Animals with unilateral or no expression of the Gi DREADD construct in cPO or BLA terminal fields were excluded from the analysis. Open arms were dimly lit (~40 lux). The time spent in each arm and the number of arm entries represented anxiety-like behavior and general locomotion, respectively. A computer equipped with MED-PC (Med Associates) connected to the plus maze collected beam breaks; and EPM test sessions were also videotaped (Panasonic Model #P0H1000, Rockville, MD) to generate heat maps of the centroid for each animal using EthoVision XT (Version 11.5, Noldus Technology, Leesburg, VA). The apparatus was cleaned with warm water/mild soap and thoroughly dried between animals.

### Statistics

All data analyses were conducted using GraphPad Prism 8.0 (GraphPad Software Inc., La Jolla, CA). Data were analyzed via t-tests or analysis of variance (ANOVA), with Bonferroni post-hoc tests used to determine the locus of effect, where appropriate. Significance was set at p<0.05. Data are presented as mean ± SEM throughout the text and figures.

## Results

### Withdrawal increases release probability at cPO inputs onto LA neurons but does not alter chemogenetic inhibition

To confirm that cPO terminals in the BLA could regulate neurophysiological changes that emerge during ethanol withdrawal, we co-injected Channelrhodopsin with the Gi-DREADD viral construct into the cPO. Expression of both the Channelrhodopsin and Gi-DREADD constructs produced intense fluorescence within the cPO (mCherry fluorescence shown in Fig. 1A_1_). Current-clamp recordings of a cPO neurons (Fig. 1B_1_) revealed that light exposures of increasing durations produced depolarizations that reached threshold at durations ≥4msec. Light-gated (45msec duration) currents measured in voltage-clamp in the presence of picrotoxin (see Methods) were insensitive to bath application of the glutamate ionotropic antagonists DNQX (20μM) and APV (50μM; Fig 1B_2_).

**Figure 1.**
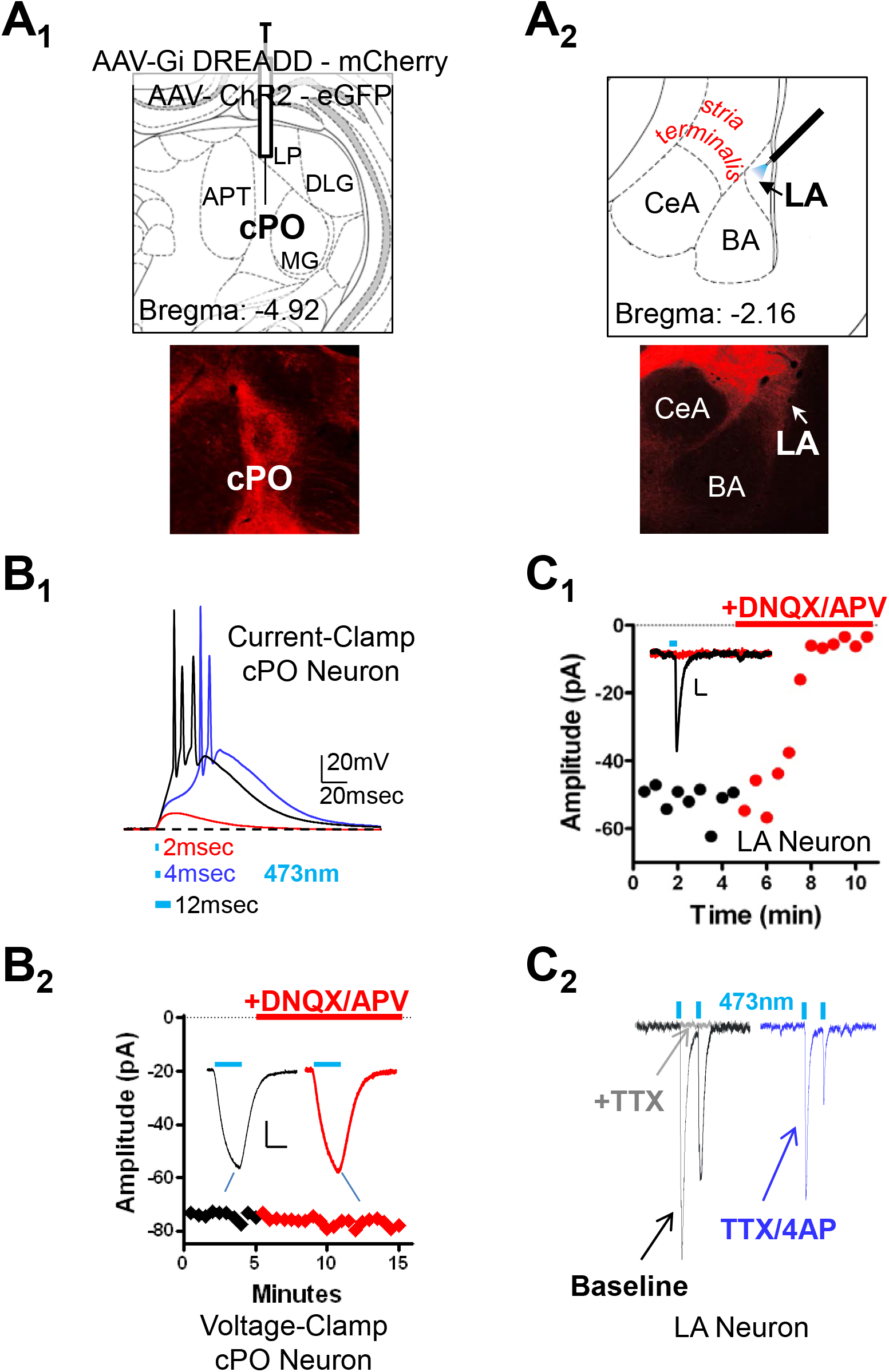
Expression of Channelrhodopsin in cPO produces monosynaptic glutamatergic responses in lateral amygdala principal neurons. (**A**) Immunofluorescence of mCherry expressed in the cPO injection site (A_1_, left) and cPO projections along the *stria terminalis* dorsal to the central amygdala (CeA) that terminate in the lateral amygdala (LA, A_2_, right). MG: medial geniculate. BA: basolateral amygdala. (**B**) Recordings from cPO neurons showing that increasing light durations cause depolarizations that transition to action potentials at stimulations >4msec (B_1_). Voltage-clamp recordings of light-gated whole-cell currents in cPO neurons were insensitive to the ionotropic glutamate receptor antagonists DNQX (AMPA receptor antagonist, 100μM) and APV (NMDA receptor antagonist, 20μM; B_2_). (**C**) Characterization of light-gated synaptic responses recorded from lateral amygdala principal neurons. Optical stimulation of cPO terminals (5msec light duration) evoked synaptic responses that were inhibited by the ionotropic glutamate receptor antagonists DNQX and APV (C_1_). These responses were monosynaptic (C_2_) as demonstrated by their complete inhibition by the voltage-gated sodium channel antagonist, tetrodotoxin (TTX, 1μM; compare baseline/black trace with TTX/gray trace) and subsequent recovery with the addition of the voltage-gated potassium channel antagonist, 4-AP (2mM, blue trace). For B & C, the width of the bright blue lines above the traces represent the duration of light stimulation (50msec in B_2_; 5msec in C). Calibration bars in B_2_ and C_1_: y = 20pA, x = 40msec.

Expression of Channelrhodopsin/Gi-DREADD also produced intense fluorescence along the stria terminalis (dorsal to the central amygdala, CeA) and within the lateral nucleus of the amygdala (LA, Fig. 1A_2_). These findings confirm previous publications with traditional retrograde tracers showing that the cPO sends strong projections to the lateral/basolateral amygdala (Doron and Ledoux, 1999; Doron and Ledoux, 2000; Lanuza et al., 2008). However, these cPO inputs appear to be largely localized in the lateral amygdala. Indeed, expression of Channelrhodopsin in cPO terminals produced light-gated synaptic responses recorded from lateral amygdala neurons that were inhibited by DNQX/APV (Fig. 1C_1_). Importantly, TTX (1μM) completely inhibited these light-evoked synaptic responses; and, bath application of the voltage-gated potassium channel blocker, 4-AP (2mM), largely reversed the inhibition by TTX (Fig. 1C_2_). Together, these data suggest that cPO neurons send monosynaptic glutamatergic projections to lateral amygdala principal neurons.

Previous work using electrical stimulation has shown that chronic ethanol exposure can increase presynaptic function of stria terminalis inputs onto lateral and basolateral amygdala neurons (Christian et al., 2013; Morales et al., 2018). We utilized a similar paired-pulse protocol that instead elicited light-evoked responses with an inter-stimulus duration of 50msec to measure the effects of chronic ethanol exposure specifically on cPO terminals onto LA neurons. We found here that CIE exposure also increases glutamatergic release probability from cPO terminals onto lateral amygdala neurons. Neurons from air-exposed controls (n = 7) expressed paired-pulse ratios of −0.16±0.03 while those from CIE/24h withdrawal animals (n = 10) had a ratio of −0.41±0.03 (p<0.05, t-test; Fig. 2A). Like findings with electrical stimulation of the stria terminalis (Christian et al., 2013; Morales et al., 2018), these data show that CIE/withdrawal increases glutamate release from cPO inputs onto LA neurons.

**Figure 2.**
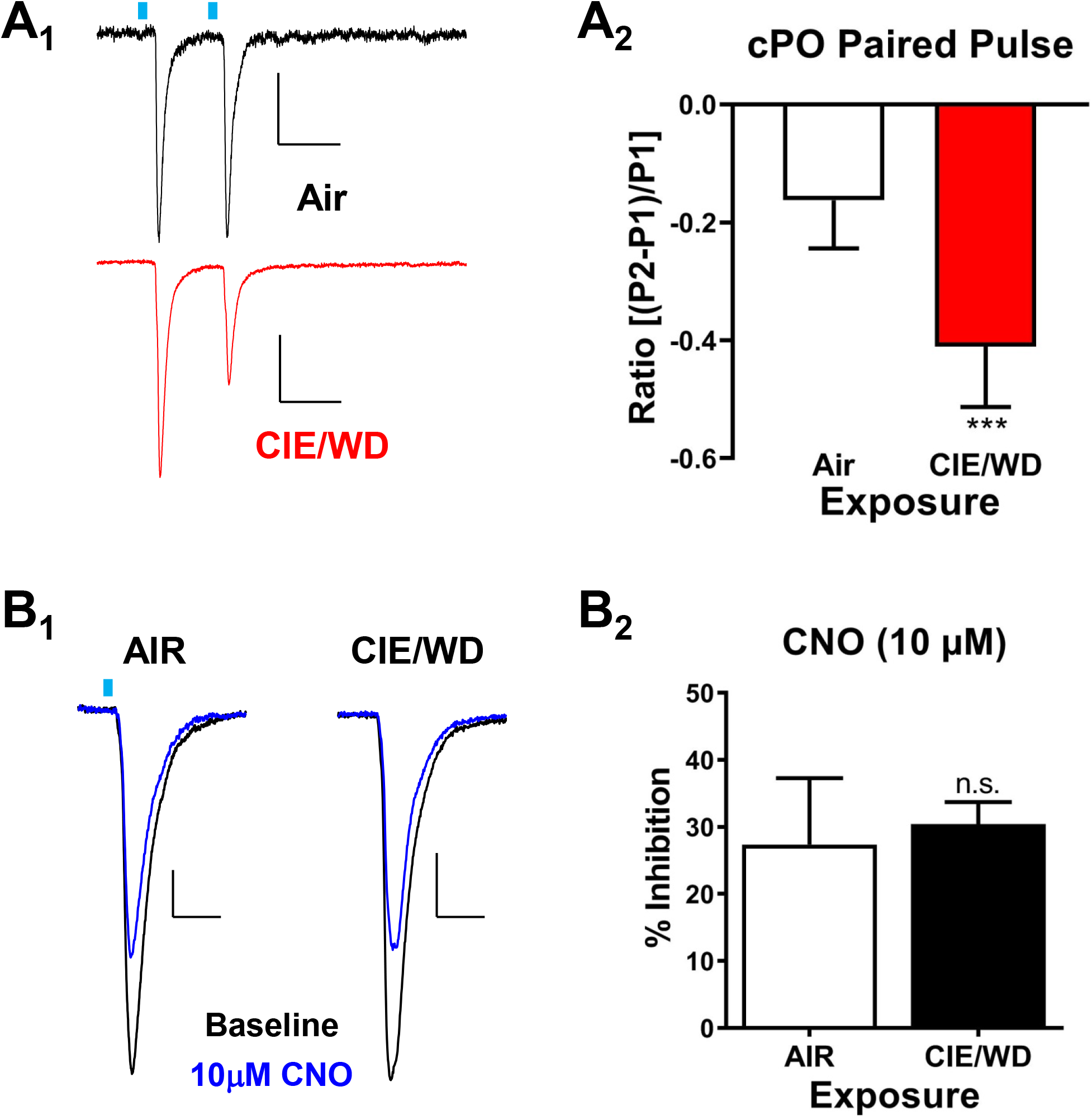
Withdrawal from chronic intermittent ethanol (CIE/WD) increases glutamate release probability at cPO-LA inputs but does not disrupt hM4-Gi-DREADD modulation of cPO terminals and bath application of CNO (10 μM) decreases glutamate release probability. (**A**) CIE/WD decreases the paired pulse ratio of light-gated responses from cPO terminals. Representative traces (A_1_) from air-exposed control neurons (top, black trace) and CIE/WD neurons (bottom, red trace) were normalized to the amplitude of the first response to emphasize the relationships with the amplitude of the second response across treatment groups. Calibration bars: x = 50msec; y = 20pA for the Air trace, 50 pA for the CIE/WD trace. Summary of paired-pulse ratio data for exposure and CNO treatment groups (A_2_) shows a significant decrease in the ratio recorded from CIE/WD neurons (n=7) relative to Air/control neurons (n=10). *** -- p<0.001, Student’s t-test. (**B**) CIE/WD does not disrupt cPO terminal modulation by the Gi-DREADD agonist, CNO (10μM). Representative traces (B_1_) illustrate the effects of CNO application in the two treatment groups. Response amplitudes have been normalized across treatment group to allow comparisons with the CNO (calibration bars: x=20msec, y = 20pA in Air neuron, 40pA in CIE/WD neuron). Summary of CNO inhibition across the treatment groups (B_2_) shows no significant alterations in hM4-Gi-DREADD modulation by the CIE/WD treatment (‘n.s.’ = not significant).

To understand the behavioral role of CIE-dependent alterations in glutamate release from cPO terminals in the LA, we examined Gi-DREADD modulation of light-evoked responses from these cPO-LA terminals from either air controls or CIE-exposed animals 24h after the last exposure (Fig. 2B_1_). Bath application of CNO (10μM) decreased glutamate release from cPO terminals from both treatment groups to a similar extent (Fig. 2B_2_). CNO inhibition was 27.4±9.9% in air-exposed neurons (n=5) and 30.4±3.3% in CIE/24h withdrawal neurons (n=6; p>0.05, t-test). CNO inhibition did not appear to reverse even after extensive wash out (10min).

### Systemic Injection of CNO Reverses Withdrawal-associated Anxiety-like Behavior

To understand the potential role of cPO in the expression of anxiety-like behavior following chronic ethanol exposure, we expressed the Gi DREADD in cPO neurons, exposed them to CIE, and measured the effects of systemically administered saline or CNO (5mg/kg, I.P.) 24h after the last ethanol exposure on the elevated plus maze (Figure 3A). Two-way ANOVA of open arm time (Figure 3B_1_) revealed a significant main effect of CNO treatment (p<0.05, F[1, 24]=6.697); and, withdrawal animals injected with CNO spent significantly more time in the open arm compared to their saline counterparts (p<0.05, Bonferroni’s multiple comparisons post-test). For open arm entries, the two-way ANOVA showed significant main-effects of both WD (F[1,24]=4.84, p<0.05) and CNO (F[1,24]=5.45, p<0.05) but no interaction (F[1,24]=3.04, p>0.05). Air+CNO animals made significantly more open arm entries than the Air+Saline group (p<0.05, Bonferroni’s multiple comparison post-test); but open arm entries was not significantly altered by CNO in the WD group (p>0.05). There were no significant interactions (F[1,24]=0.2.459, p>0.05) or main effect of CNO (F[1,25]=2.768, p>0.05) between treatment groups for total arm entries (Figure 3B_2_) or closed arm entries (interaction – F[1,25]=0.001, p>0.05, WD – F[1,25]=1.19, p>0.05, CNO – F[1,25]=0.01, p>0.05). There was a significant main effect of WD on total arm entries (F[1,25]=7.579, p<0.05) that appears to be driven by the impact of WD on open arm entries. These data all suggest that there was no substantial locomotor effects of the chronic ethanol exposure or the CNO.

**Figure 3.**
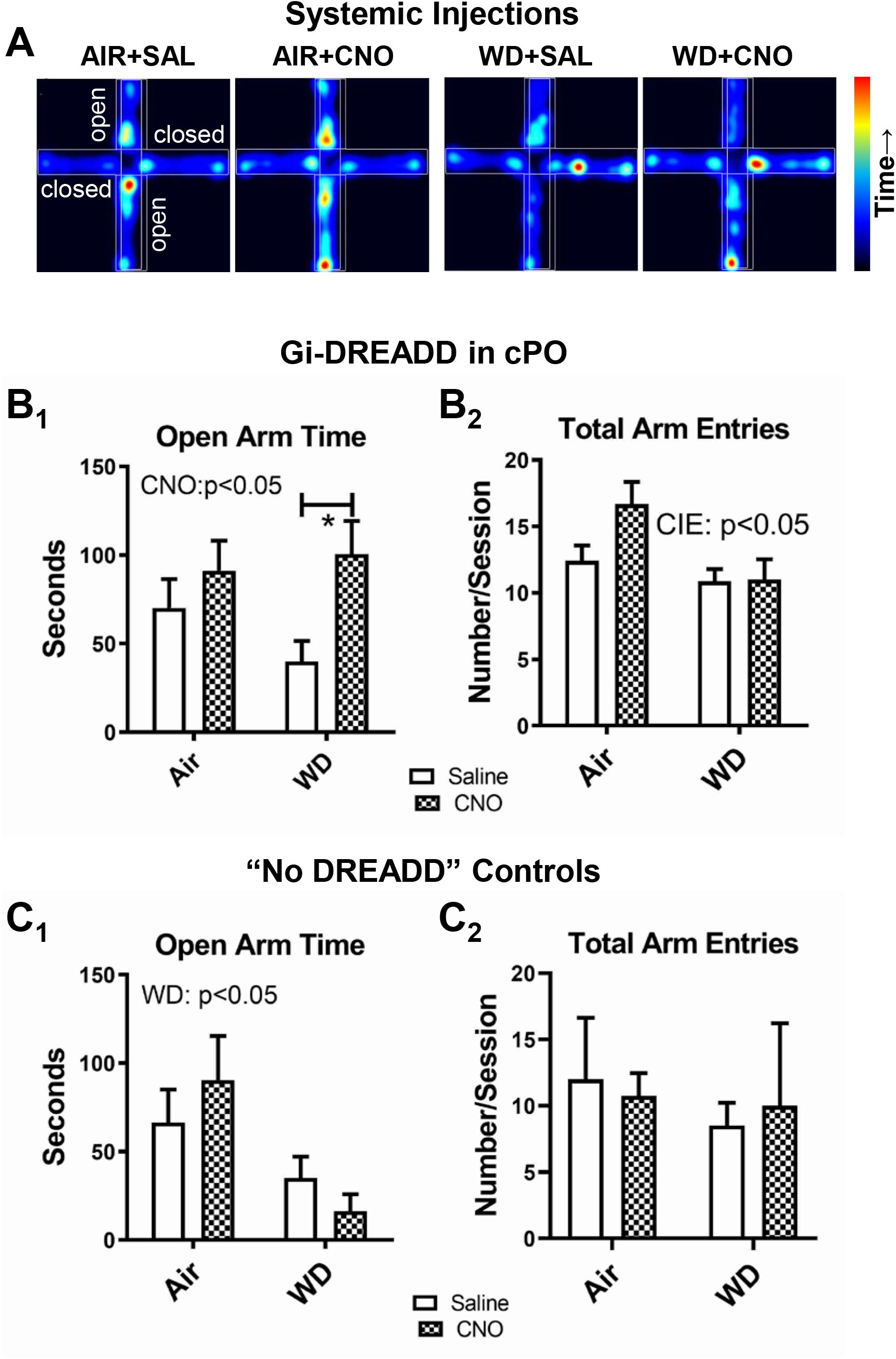
Systemic activation of hM4-Gi-DREADD expressed in the cPO attenuates anxiety-like behavior expressed during withdrawal. (**A**) Representative heat maps demonstrating location/duration of time spent on the open and closed arms for each group (Air controls ± CNO, CIE/WD ± CNO). Calibration bar (right) indicates color code for time duration in each location. (**B**) Intraperitoneal injection of CNO (5 mg/kg) significantly increased open arm time (B_1_) relative to saline (1ml/kg) in CIE/WD animals (n=8 CIE/WD+saline, n=7 CIE/WD+CNO) but not air-exposed controls (n=7 Air+saline, n=6 Air+CNO). There was a significant main effect of CNO (p<0.05, two-way ANOVA) with a significant increase in open arm time in the CIE/WD+CNO group compared to the CIE/WD+saline group (* - p<0.05, Bonferroni’s multiple comparison test). As a proxy for locomotor behavior, there was no significant effect of CNO on closed arm entries in either treatment group (B_2_). (**C**) Systemic CNO modulation of anxiety-like behavior requires expression of the hM4-Gi-DREADD in the cPO. In animals expressing only Channelrhodopsin in the cPO (“No DREADD” Controls), there was no significant effect of CNO systemic injection on either open arm time (C_1_) or closed arm entries (C_2_). However, CIE/WD animals (n=4, CIE/WD+saline, n=4, CIE/WD+CNO) spent significantly less time in the open arms compared to Air-exposed controls (n=5, Air+saline, n=4 Air+CNO; main effect of CIE/WD, p<0.05, two-way ANOVA).

To confirm that the observed CNO effects were specific to cPO neurons expressing the DREADD receptor, a separate group of rats underwent surgery but were only injected with Channelrhodopsin. Using the same experimental design, rats from air- or CIE-treatments were injected with either saline or CNO (5 mg/kg, I.P.) 30min prior to measurement on the elevated plus maze. The two-way ANOVA of the data from these “no DREADD” controls showed a significant main effect of WD on open-arm time (Figure 3C_1_; F[1,13]=8.37, p<0.05) compared to the Air controls but no interaction (F[1,13]=1.459, p>0.05) and no main effect of CNO (F[1,13]=0.02, p>0.05). Unlike the DREADD-expressing animals, open arm entries were not significantly different for either the WD (F[1,13]=3.186, p>0.05) or CNO factors (F[1,13]=0.71, p>0.05); and there was no interaction between factors (F[1,13]=1.18, p>0.05). There was again no significant interactions between the ethanol exposure and CNO injection (F[1,13]=0.471, p>0.05) and main effects of WD (F[1,13]=1.125, p>0.05) or CNO (F[1,13]=0.003, p>0.05) on total arm entries (Figure 3C_2_) or closed arm entries (interaction – F[1,13]=0.02, p>0.05; WD – F[1,13]=0.09, p>0.05; CNO – F[1,13]=0.581, p>0.05).

### Gi DREADD inhibition of cPO terminals in the Lateral Amygdala Attenuates Withdrawal-induced Anxiety-like Behavior

Results from our systemic CNO injections strongly suggest a role for the cPO in modulating anxiety-like behavior following ethanol withdrawal. We directly examined the contributions of cPO terminals in the lateral amygdala by expressing the hM4D Gi-DREADD in cPO, microinjecting the agonist CNO into the LA to inhibit glutamate release from these terminals, and measuring anxiety-like behavior on the elevated plus maze following air or CIE exposure (Fig. 4A & B). For open arm time (Fig. 4C_1_), there was a significant interaction (F[1,22]=4.88, p<0.05) between treatment (Air vs. CIE/WD) and microinjection (Vehicle vs. CNO) groups. Post-test analysis showed that this was due to a significant increase in open arm time following CNO microinjection in the CIE/WD group (p<0.05, Bonferroni’s multiple comparisons test) but not the Air group. There were no significant effects of either CIE/WD or CNO microinjection for the closed arm time, junction time, or open arm entry variables (p>0.05, two-way ANOVA). Total arm entries (Fig. 4C_2_), a measure of locomotor activity in the plus maze, were significantly less in the CIE-treated groups (main effect, F[1,22]=8.771, p<0.01) but there was no main effect of CNO microinjection (F[1,22]=0.01, p>0.05) and no interaction between CIE treatment and CNO microinjection (F[1,22]=0.60, p>0.05). Thus, while CIE had an effect on locomotor behavior in this experiment, CNO microinjection did not. This contrasts with the significant effect of CNO microinjection on open arm time in the CIE-treated animals but not air controls.

**Figure 4.**
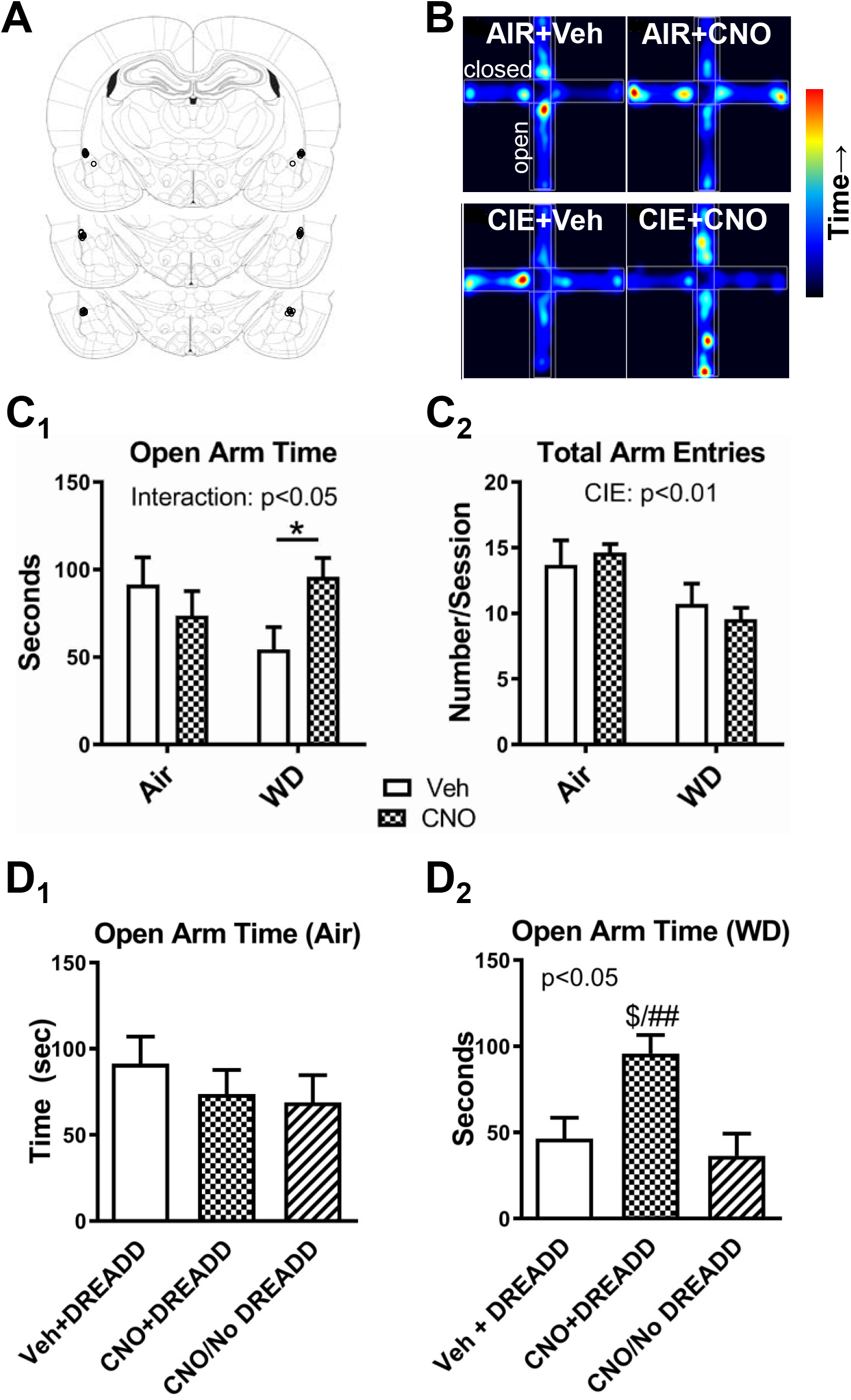
Activation of hM4-Gi-DREADD receptors expressed cPO terminals in the lateral amygdala attenuates withdrawal-dependent anxiety-like behavior on the elevated plus maze. (**A**) Illustration of the approximate placement of guide cannulae (open circles) used in this analysis. Coronal brain slices have been modified from Paxinos and Watson (Paxinos and Watson, 2005). (**B**) Representative heat maps from the four treatment groups. (**C**) Microinjection of CNO into the lateral amygdala decreases anxiety-like behavior in CIE/WD animals but not air controls. There was a significant interaction between treatments (Air vs. CIE/WD and vehicle vs. CNO, two-way ANOVA) and a significant increase in open arm time in CIE/WD animals microinjected with CNO (*, p<0.05, Bonferonni’s multiple comparison post-test). There was a significant main effect of CIE/WD on total arm entries (C2) but no effect of CNO microinjection. (**D**) Modulation of withdrawal-associated anxiety-like behavior by CNO microinjection requires cPO expression of the hM4-Gi-DREADD. For the air-treated control groups (D_1_), there was no significant difference between vehicle microinjection in Gi-DREADD animals, CNO microinjection in Gi-DREADD animals (data from Fig. 4C), and CNO microinjection in animals expressing only Channelrhodopsin (“No DREAD” controls, n=4; p>0.05, on-way ANOVA). In the CIE/WD groups (D_2_), attenuation of anxiety-like behavior required both Gi-DREADD expression in cPO terminals and CNO microinjection into the lateral amgydala (p<0.05, one-way ANOVA; $ - p<0.05 relative to vehicle injected animals, ## - p<0.01 relative to “No DREADD” controls, n=5, Bonferroni’s multiple comparison test).

Like the systemic studies, we also examined the effects of CNO microinjection into the LA following air (n=4) or CIE treatment (n=5) in animals expressing ‘empty’ (no DREADD) vector in cPO terminals in a separate group of animals. Direct comparisons with the air and CIE groups from the previous experiment revealed that CNO had no effect on open arm time when the Gi DREADD was absent from cPO terminals (Fig. 4D_1_ & D_2_). Separate analysis of the air exposed animals showed no significant differences between the DREADD+Veh, DREADD+CNO, and No DREADD+CNO groups (F[2,11]=0.60, p>0.05). In the CIE-treated group, there was a significant difference between injection groups (F[2,18]=6.453, p<0.01); and, CNO microinjection significantly increased open arm time in the DREADD+CNO animals relative to both the DREADD+Veh (p<0.05, Bonferroni’s multiple comparisons test) and No DREADD + CNO (p<0.01) rats. There were no significant differences between the DREADD+Veh, DREADD+CNO, and No DREADD+CNO microinjections for closed arm time (Air – F[2,11]=0.816, p>0.05; CIE – F[2,18]=0.327, p>0.05) or for total arm entries (Air – F[2,11]=0.762, p>0.05; CIE – F[2,18]=3.195, p>0.05). These findings together suggest that the increased open arm time caused by microinjection of CNO requires DREADD expression on cPO terminals in the LA.

## Discussion

In the current work we showed that microinjection of Channelrhodopsin into the cPO neurons labels terminal fields in the lateral amygdala and that these inputs are both glutamatergic and monosynaptic onto LA principal neurons. This is consistent with previous work showing that the posterior intralaminar and subgeniculate thalamic nuclei provide robust projections to the lateral amygdala (Doron and Ledoux, 1999; Gauriau and Bernard, 2004; Linke et al., 2000; Smith et al., 2019), can drive local field potentials in an *in vivo* preparation (Doyere et al., 2003), and are glutamatergic (LeDoux and Farb, 1991). Withdrawal from chronic ethanol inhalation enhances light-evoked glutamate release probability from these synapses similar to previous findings with electrical stimulation showing that this ethanol exposure increases release from *stria terminalis* inputs to the lateral/basolateral amygdala (Christian et al., 2013; Morales et al., 2018). Co-expression of the hM4-Gi-DREADD in cPO terminals conferred inhibition by the DREADD agonist, CNO, that was not impacted by chronic ethanol at the concentration used here (10μM). Furthermore, systemic injection of CNO reduced anxiety-like behavior on the elevated plus maze in CIE-exposed rats, but not air-treated controls, when those animals expressed the Gi-DREADD in the cPO. Local activation of Gi DREADDS expressed on cPO terminals in the BLA using microinjection also attenuated withdrawal-anxiety. Finally, both the systemic effects of CNO and the effects of CNO microinjection were absent when Gi-DREADDs were not expressed indicating the effects of the agonist were DREADD-specific. Together, our findings suggest chronic ethanol/withdrawal facilitates the function of cPO terminals in the lateral amygdala and that this facilitation helps increase the expression of anxiety-like behaviors during withdrawal.

Our findings are consistent with previous literature showing that sub-regions within the cPO help regulate aversive behaviors by providing somatosensory input to the lateral amygdala. These behaviors can include unconditioned responses to itch/nociceptive (Gauriau and Bernard, 2004; Lipshetz et al., 2018), tactile (Gauriau and Bernard, 2004; Gonzalez-Hernandez et al., 2013), auditory (Bordi and LeDoux, 1994a; Bordi and LeDoux, 1994b), and visual stimuli (Eordegh et al., 2005). The cPO also regulates conditioned responses to somatosensory stimuli (Halverson et al., 2008; Romanski and LeDoux, 1992). Importantly, recent work using optogenetic approaches showed that cPO inputs to the lateral amygdala play a substantial role in the acquisition of these conditioned behaviors and that this process is blocked by local (LA) microinjection of ionotropic glutamate receptor antagonists (Kwon et al., 2014). These studies highlight the importance of cPO-LA circuitry in the expression of aversive behaviors.

Recent studies also suggest a strong relationship between the cPO-LA circuit and reward-related behaviors. For example, natural rewards like social play and food increase the expression of ‘neuronal activity’ markers like cFos (van Kerkhof et al., 2014) and phosphoMAP kinase (Nasser and McNally, 2013) in the lateral amygdala. Most notably, the acquisition of cue-sucrose associations increased LA neuron firing in response to cue when measured with *in vivo* preparations and increased AMPA receptor synaptic function at *stria terminalis* inputs (but not external capsule); this increased postsynaptic function was positively correlated with task efficiency and accuracy (Tye et al., 2008). Similar findings for drug-related rewards like cocaine are also in the literature – from increased expression of markers for neuronal activity (Thomas and Everitt, 2001) to increased synaptic function at *stria terminalis* inputs to LA neurons (Goussakov et al., 2006). In a recent study, Rich and colleagues (Rich et al., 2019) examined the effects of cue-cocaine pairings, extinction, and reinstatement of LA neuron synaptic function. This study found bi-directional modulation of *stria terminalis* synapses onto LA associated with both learned cue-cocaine associations (facilitation) and extinction (depression). This study also utilized optogenetic approaches that expressed Channelrhodopsin around the medial geniculate (including most of the cPO) and found the same bidirectional shifts in light-gated EPSCs from these terminals recorded from LA neurons. These studies, together with the current work, highlight the important role of cPO-lateral amygdala circuits integrating somatosensory information with emotionally relevant behaviors.

Finally, it is worth noting that the non-contingent ethanol exposure used in the current work produces presynaptic facilitation of glutamate release at cPO inputs to the LA. The presynaptic facilitation described here using optogenetics is remarkably similar to previous work with electrical stimulation (Christian et al., 2013; Lack et al., 2009; Lack et al., 2007; Morales et al., 2018). In contrast, studies using contingent delivery of natural or drug rewards paired with cues have either suggested or directly demonstrated postsynaptic facilitation of synaptic function at these same inputs (Rich et al., 2019; Tye et al., 2008). Neither the Tye et al. study nor the Rich et al. study, which used both electrical stimulation of the stria terminalis as well as optogenetic activation of thalamic terminals, found any effects of the conditioning on presynaptic function of thalamic input pathways. Results from aversive-conditioning studies are more mixed with some showing that cue-aversion associations produce post-synaptic facilitation (Kim et al., 2007; Park et al., 2016) while others showing increased presynaptic function at *stria terminalis* inputs (Park et al., 2016; Shinnick-Gallagher et al., 2003). Interestingly, using auditory fear conditioning, Park and colleagues (Park et al., 2016) showed that the synaptic locus of facilitation (pre-versus post-) appears to depend upon the complexity of the tone used in the conditioning paradigm with ‘less-complex’ auditory stimuli producing greater presynaptic facilitation. The *stria terminalis* inputs thus appear to have the capacity to express both pre- and postsynaptic plasticity. This has been demonstrated directly using *in vitro* preparations. Shin et al. (Shin et al., 2010) showed that presynaptic ‘plasticity’ of *stria* inputs requires the activation of presynaptic kainite receptors expressed on these terminals. In contrast, pairing postsynaptic depolarization with activation of the same *stria* projections elicits postsynaptic facilitation that subsequently occludes any further changes in presynaptic function via the release of endogenous endocannabinoids and activation of presynaptic CB1 receptors expressed on these terminals. It is noteworthy then that the acquisition of conditioned behaviors is associated with increased firing of LA neurons in response to cue presentation (Tye et al., 2008). Regardless, it is not yet clear whether presynaptic facilitation by chronic ethanol at *stria terminalis* inputs is specific to ethanol or are related to either the contingency of drug delivery or the severity of intoxication during the exposure. But, recent work from our lab has shown that acute ethanol can directly inhibit presynaptic function of glutamate synapses during high frequency activity (Gioia et al., 2017; Gioia and McCool, 2017). This suggests that presynaptic facilitation following chronic ethanol exposure may represent a compensatory up-regulation in response to this acute suppression.

In conclusion, using optogenetic approaches, we have shown that withdrawal from chronic ethanol facilitates the presynaptic function of somatosensory thalamic inputs to lateral amygdala principal neurons. Subsequent down-regulation of these cPO terminals with chemogenetic approaches was sufficient to reduce the expression of withdrawal-related anxiety-like behaviors. These data highlight the role of circuits integrating somatosensory perception with emotional behaviors following chronic ethanol exposure. Future experiments should explore the relationship of synaptic function and locus of adaptation along with contingent delivery of ethanol and severity of intoxication.

Author Contributions
MM and BAM designed research; MM, MMM, AEC, BPP, and BAM performed experiments; MM, MMM, and BAM analyzed data; MM and BAM wrote the manuscript

## Acknowledgements

This work was supported by the following grants from the National Institutes of Health/National Institute on Alcohol Abuse and Alcoholism – R01 AA014445 (BAM), R01 AA023999, P50 AA026116 (BAM), R21 AA026572 (BAM), T32 AA007565 (MM, MMM), F32 AA024949 (MM), and F31 AA025514 (MMM)

